# Hepatitis B virus Core protein nuclear interactome identifies SRSF10 as a host RNA-binding protein restricting HBV RNA production

**DOI:** 10.1101/2020.05.04.076646

**Authors:** Hélène Chabrolles, Héloïse Auclair, Serena Vegna, Thomas Lahlali, Caroline Pons, Maud Michelet, Yohann Couté, Lucid Belmudes, Yujin Kim, Ariel Di Bernardo, Pascal Jalaguier, Michel Rivoire, Lee D. Arnold, Uri Lopatin, Christophe Combet, Fabien Zoulim, David Grierson, Benoit Chabot, Julie Lucifora, David Durantel, Anna Salvetti

## Abstract

Despite the existence of a preventive vaccine, chronic infection with Hepatitis B virus (HBV) affects more than 250 million people and represents a major global cause of hepatocellular carcinoma (HCC) worldwide. Current clinical treatments, in most of cases, do not eliminate viral genome that persists as a DNA episome in the nucleus of hepatocytes and constitutes a stable template for the continuous expression of viral genes. Several studies suggest that, among viral factors, the HBV core protein (HBc), well-known for its structural role in the cytoplasm, could have critical regulatory functions in the nucleus of infected hepatocytes. To elucidate these functions, we performed a proteomic analysis of HBc-interacting host-factors in the nucleus of differentiated human hepatocytes. The HBc interactome was found to consist primarily of RNA-binding proteins (RBPs), which are involved in various aspects of mRNA metabolism. Among them, we focused our studies on SRSF10, a RBP that was previously shown to regulate alternative splicing in a phosphorylation-dependent manner and to control stress and DNA damage responses, as well as viral replication. Functional studies combining SRSF10 knockdown and a pharmacological inhibitor of SRSF10 phosphorylation (1C8) showed that SRSF10 behaves as a restriction factor that regulates HBV RNAs levels and that its dephosphorylated form is likely responsible for the anti-viral effect. Surprisingly, neither SRSF10 knock-down nor 1C8 treatment modified the splicing of HBV RNAs but rather modulated the level of nascent HBV RNA. Altogether, our work suggests that in the nucleus of infected cells HBc interacts with multiple RBPs that regulate viral RNA metabolism. Our identification of SRSF10 as a new anti-HBV restriction factor offers new perspectives for the development of new host-targeted antiviral strategies.

**Author Summary:** Chronic infection with Hepatitis B virus (HBV) affects more than 250 millions of people world-wide and is a major global cause of liver cancer. Current treatments lead to a significant reduction of viremia in patients. However, viral clearance is rarely obtained and the persistence of the HBV genome in the hepatocyte’s nucleus generates a stable source of viral RNAs and subsequently proteins which play important roles in immune escape mechanisms and liver disease progression. Therapies aiming at efficiently and durably eliminating viral gene expression are still required. In this study, we identified the nuclear partners of the HBV Core protein (HBc) to understand how this structural protein, responsible for capsid assembly in the cytoplasm, could also regulate viral gene expression. The HBc interactome was found to consist primarily of RNA-binding proteins (RBPs). One of these RBPs, SRSF10, was demonstrated to restrict HBV RNA levels and a drug, able to alter its phosphorylation, behaved as an antiviral compound capable of reducing viral gene expression. Altogether, this study sheds new light novel regulatory functions of HBc and provides information relevant for the development of antiviral strategies aiming at preventing viral gene expression.

## Introduction

Despite the existence of a preventive vaccine, chronic infection with Hepatitis B virus (HBV) remains a major health problem worldwide, as it represents a major global cause of hepatocellular carcinoma (HCC) [1]. Clinically approved treatments, mainly based on nucleoside analogs (NUCs), can reduce HBV viremia under the limit of detection in patients [2]. NUCs, while potent, only affect a relatively late step in the viral life cycle, the conversion of viral pre-genomic RNA into viral DNA after encapsidation. They have no known effect elsewhere in the viral life cycle, and as a result viral clearance is rarely obtained and rebound off therapy is common, thus making life-long therapy with NUCs mandatory. The persistence of the viral genome (an episome called covalently-closed-circular dsDNA or cccDNA) in the nucleus of non-dividing hepatocytes constitutes one major obstacle toward a complete eradication of HBV infection. Indeed, cccDNA not only guarantees viral persistence in the organism but also constitutes a stable source of viral protein expression, including the HBe and HBs antigens (HBeAg and HBsAg), which play important roles in immune escape mechanisms and liver disease progression [3]. Therefore, therapies aiming at efficiently and durably blocking the production of viral antigens are still required [4, 5].

HBV is a small enveloped, DNA virus that replicates in hepatocytes. After binding to its receptor, the sodium taurocholate co-transporting polypeptide (NTCP), and uncoating, the viral capsid is transported to the nucleus where the viral genome, constituted by a relaxed circular and partially dsDNA molecule of 3.2 Kb (rcDNA), is released [6]. Conversion of rcDNA into cccDNA occurs in the nucleoplasm *via* the intervention of cellular enzyme [7–9]. It results in the establishment of a viral episome that constitutes the template for the transcription of five RNAs of 3.5 (precore and pregenomic RNA), 2.4, 2.1 and 0.7 kb that, respectively, encode the HBeAg, Core protein (HBc), viral polymerase, three surface glycoproteins (S, M and L; all defining the HBsAg), and X protein (HBx). Importantly, all these RNAs are unspliced. Several other spliced RNA species are also generated. These spliced forms can be detected in the sera and livers of chronically-infected patients as well as in cells transfected with HBV genomes [10]. They are not required for virus replication but could be involved in HBV-induced pathogenesis and disease progression [11]. Formation of new viral particles initiates in the cytoplasm by packaging of the polymerase-bound pregenomic RNA (pgRNA) into the capsid. Reverse transcription of pgRNA into rcDNA occurs within capsids that are then either enveloped and secreted to form progeny viral particles or re-routed toward the nucleus to replenish the cccDNA pool [6].

HBc is the sole structural component required for the assembly of the capsid [12]. This protein of 183 amino acids (aa) is composed of a N-terminal domain (NTD, aa 1-140) that is essential for the assembly process, and a C-terminal basic domain (CTD, aa 150-183) that is dispensable for assembly. The CTD domain contains motifs responsible for trafficking of the capsid in and out of the nucleus and displays DNA/RNA binding and chaperone activities [13, 14]. Studies on HBc assembly have shown that the protein forms homodimers. Capsid assembly is initiated by the slow assembly of a trimer of dimers to which HBc dimers rapidly associate to form an icosahedral capsid [12]. Packaging of Pol-pgRNA complex that occurs during capsid assembly is mediated by the CTD of HBc, which also regulates reverse-transcription of pgRNA into rcDNA [15–17].

Converging observations suggest that, besides its structural role in the cytoplasm, HBc may also exhibit important regulatory activities to control the establishment and persistence of HBV infection. First, following viral entry, HBc, derived from incoming particles, can enter the nucleus together with rcDNA, where it can form dimers/oligomers and also reassemble into “capsid-like” structures [18, 19]. Nuclear entry of HBc can occur after a *de novo* infection, or as a consequence of the re-routing of capsids to the nucleus. Accordingly, nuclear HBc can easily be detected either *in vitro*, *i.e.* in experimentally infected human hepatocytes, or *in vivo* in the livers of chronically infected patients or model animals [20–23]. Second, earlier studies have shown that HBc binds to cccDNA, and modifies nucleosomal spacing [24, 25]. Association of HBc to cccDNA was further confirmed *in vitro* and *in vivo* and correlated to an active transcriptional state [26–28]. Finally, HBc was also reported to bind to the promoter region of several cellular genes [29]. Altogether, these data strongly suggest that this structural protein may be important at some nuclear steps of the viral life cycle that remain to be clarified. In order to gain insight into HBc nuclear functions, we performed a proteomic analysis of its cellular partners in the nucleus of human hepatocytes. Our results revealed that HBc mainly interacts with a network RNA-binding proteins (RBPs) that are involved in several post-transcriptional processes and in particular, pre-mRNA splicing. Among these RBPs, we identified SRSF10 as a host factor restricting HBV RNA synthesis/accumulation which opens new perspectives for the development of novel antiviral agents.

## RESULTS

### Host RNA-binding proteins are major HBc interacting factors in the nucleus of differentiated hepatocytes

To gain insight on HBc regulatory functions, we sought to identify its nuclear host-partners in human hepatocytes. To this end, we used differentiated HepaRG cells (dHepaRG) expressing HBc, fused at its N-terminus to a streptavidin (ST)-binding peptide (dHepaRG-TR-ST-HBc) under the control of a tetracyclin-inducible promoter (Fig 1A). The ST-HBc fusion protein localized in the nucleus of hepatocytes (S1A Fig) and assembled into capsid-like structures as wild type (wt) HBc, indicating that tag addition at its N-terminus did not alter these functions (S1B Fig). ST-HBc/host-factor complexes were purified from nuclear extracts on Strep-Tactin affinity columns (Fig 1B and Fig 1C). The negative control was provided by dHepaRG-TR cells expressing wt HBc, without any tag and thus unable to bind to the affinity column. In addition, to eliminate cellular partners recovered *via* DNA/RNA bridging, purification of ST-HBc-complexes was also performed on cell lysates submitted to nucleic acid digestion with Benzonase. Three independent purifications of ST-HBc-associated proteins, done with three different HepaRG differentiation batches, were performed in each condition (+/− Benzonase) and eluted proteins were analyzed by mass spectrometry (MS)-based label-free quantitative proteomics. This analysis resulted in the identification of 60 and 45 proteins found significantly associated with HBc, with and without Benzonase treatment, respectively (p-value<0.01 and fold change>4) (S1 Table). Importantly, 38 of these factors were common to both conditions, demonstrating the reliability of their identification (Fig. 1D)

**Fig 1.**
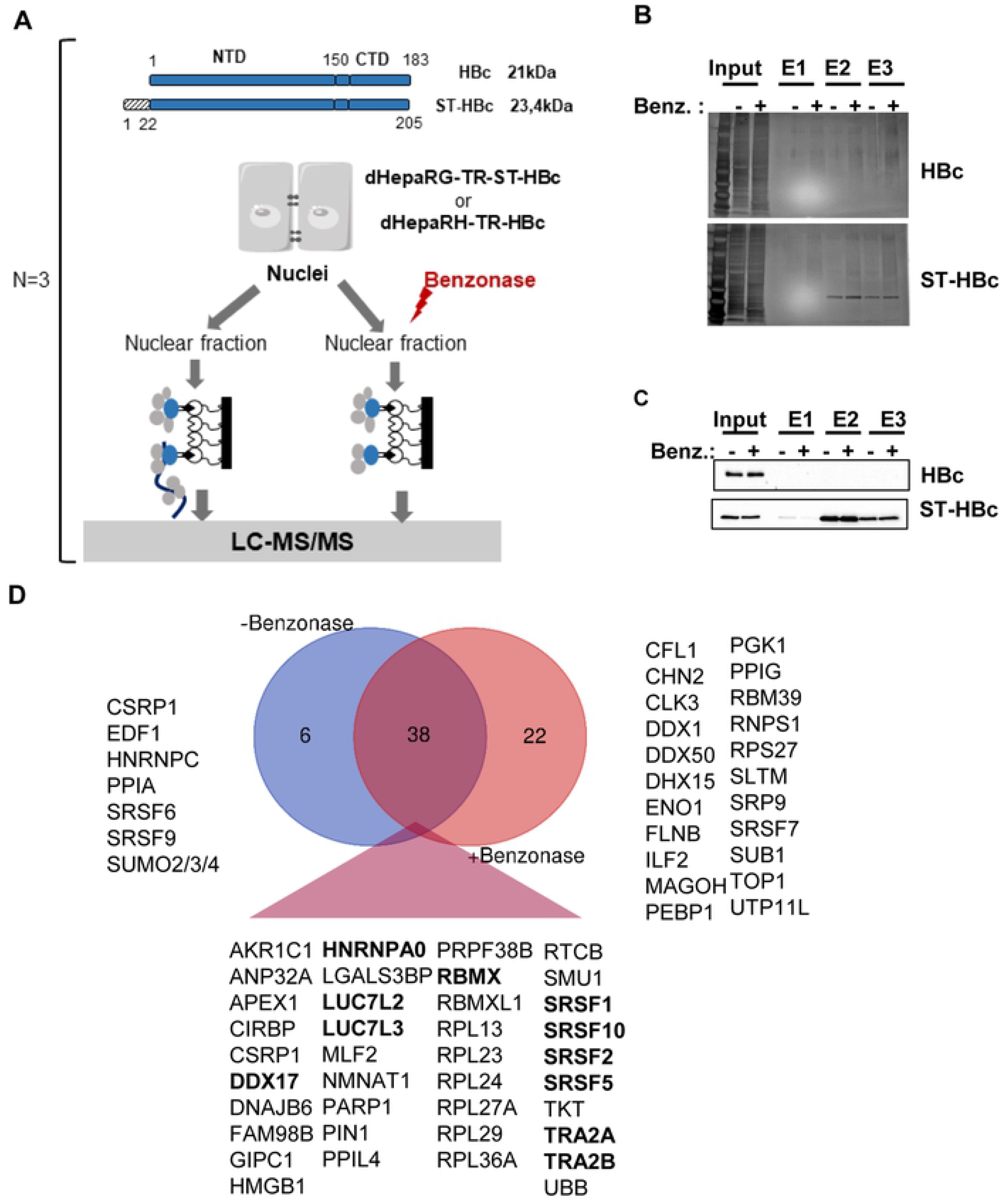
Identification of HBc-interacting proteins in the nucleus of dHepaRG cells. (A) Schematic view of HBc purification process. Nuclei were purified from differentiated HepaRG-TR cells (dHepaRG-TR) expressing either wt HBc or ST-HBc under the control of a tetracyclin-inducible promoter, lysed and then treated or not with Benzonase. Nuclear extracts were purified on a Streptactin column and protein eluted with desthiobiotin. Input and eluted fractions (E1, E2, and E3) were analyzed by gel electrophoresis followed by silver staining (B) and western blot (C) using an anti-HBc antibody. (D) Venn diagram of proteins significantly associated to HBc common to conditions with and without Benzonase. Proteins in bold correspond to the 11 “founders” RBP common to both conditions (see text, RBMXL is not highlighted because it was considered as a retrogene of RBMX).

Gene ontology (GO) annotation of HBc-interacting factors, revealed that approximately 50% of the factors, identified with or without Benzonase treatment and significantly associated with HBc, were nucleic acid binding proteins and belonged to the RBP family. In the presence of Benzonase, the most abundant protein category (Q-value: 1.8 ×10^−29^) identified, corresponded to factors involved in RNA post-transcriptional processes, in particular splicing (Fig 2A). The second most-relevant category (Q-value: 4.4×10^−14^) corresponded to ribosomal proteins. The same functional categories were retrieved using the list of proteins significantly associated to HBc without Benzonase treatment (data not shown).

**Fig 2.**
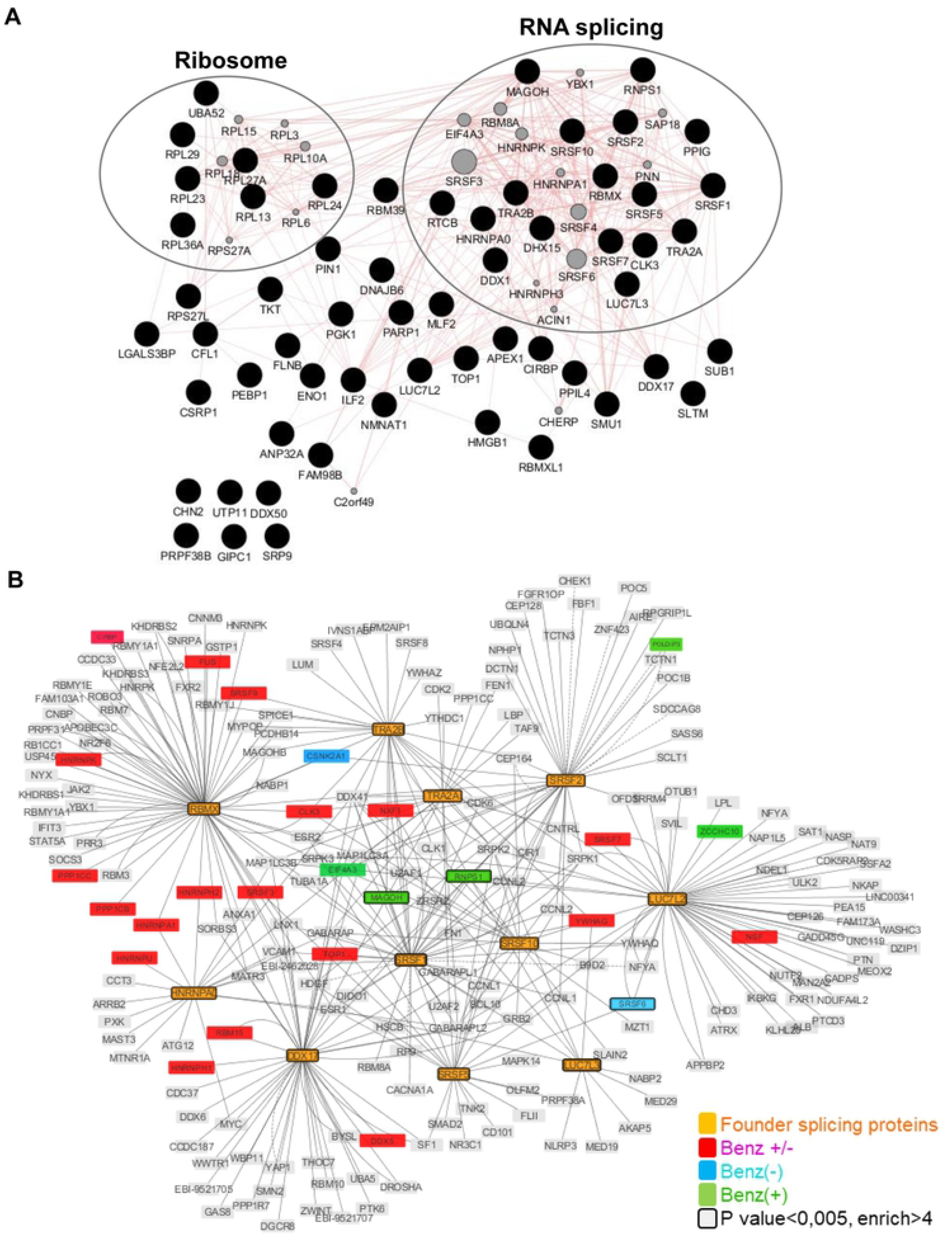
HBc nuclear interactome. (A) Proteins significantly associated to HBc in the presence of Benzonase were analyzed using the Genemania plugin in Cytoscape (3.7.1). Red lines indicate known physical interactions. Missing nodes are indicated by grey circles. (B) **Interaction network of proteins involved in mRNA splicing via spliceosome.** Significant proteins, common to the Benzonase−/+ conditions, over-representing the mRNA splicing via spliceosome biological process (“founder proteins” highlighted as orange nodes) were used to initiate the network by querying IntAct database. Red, blue and green nodes denote protein of the computed network that are found in the proteomic hits of both Benzonase−/+ (Benz− or Benz+) conditions. Nodes with a bold border indicate significant proteins (p-value<0.005 and fold change>4) from the proteomics data.

As the major GO category corresponded to RBPs involved in splicing, we next focused on proteins corresponding to this functional group and common to conditions with and without Benzonase (*i.e.* 11 proteins highlighted in bold in figure 1D). The interactome of these 11 RBPs, hereafter designed as “founder” RBPs, showed that they were highly inter-connected and that several of their first-level interacting partners were also found among HBc-co-purified factors (Fig 2B).

The analysis of the relative abundance of these founder RBPs indicated that SRSF10 was the most abundant RBPs co-purified in HBc-complexes, followed by RBMX, SRSF1, SRSF5 and TRA2B (Fig 3A). Western blot analyses confirmed the presence of SRSF10, RBMX, DDX17, SRSF2 and TRA2B in ST-HBc purified complexes, as well as that of two other non-RBP factors, PARP1 and DNAJB2 (Fig 3B-C). In contrast, the presence of SRSF1 could not be confirmed by Western blot (Fig 3B). The reason for this lack of detection is presently unclear but it could be due to a poor sensitivity of the antibodies used. The interaction between HBc and SRSF10 was confirmed by co-immunoprecipitation (co-IP) analyses performed in dHepaRG cells expressing wt HBc, indicating that this interaction occurred independently of the tag (Fig 3D).

**Fig 3.**
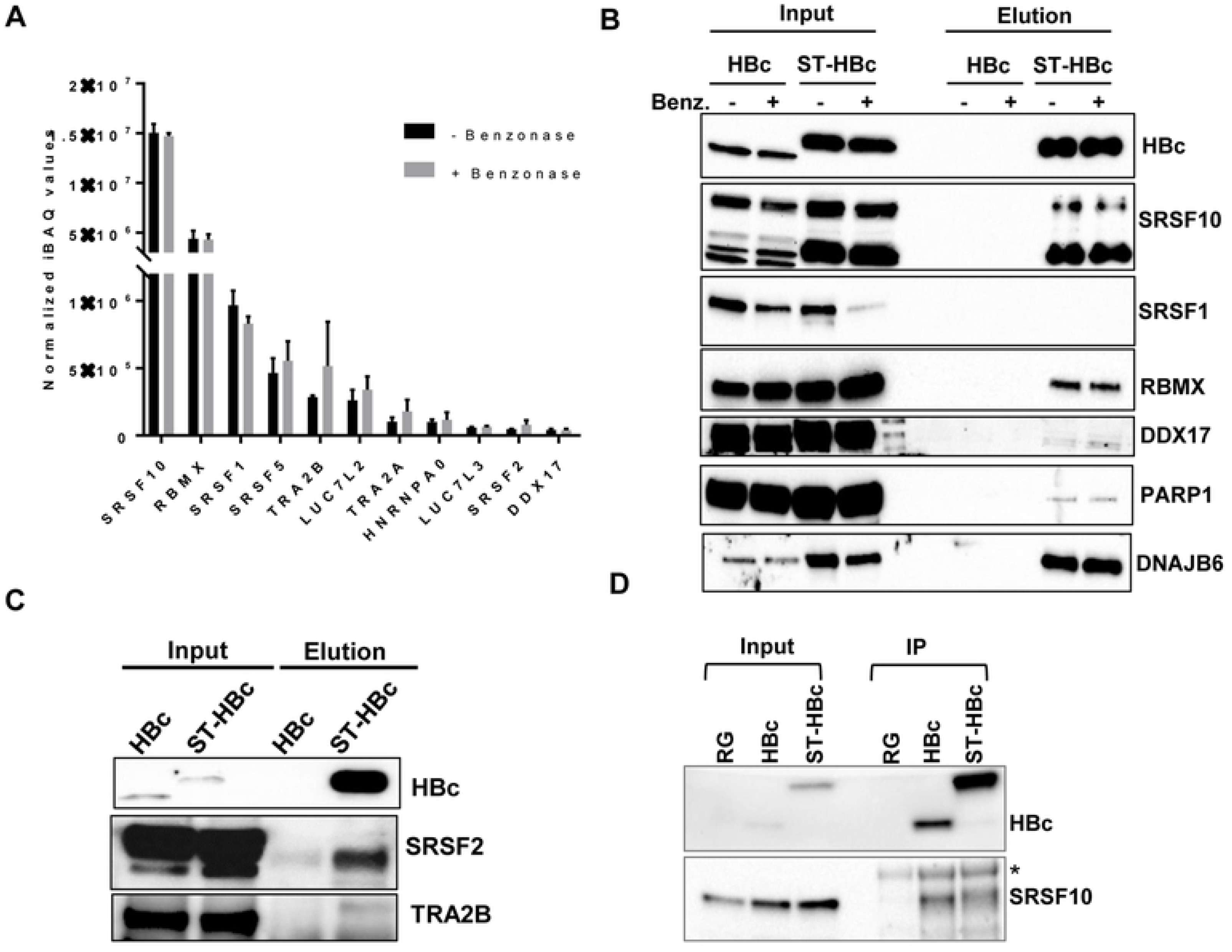
Validation analyses. (A) Relative abundances of the 11”founder” RBPs identified in HBc nuclear complexes submitted or not to Benzonase treatment. The relative abundances of HBc binding partners have been evaluated using the iBAQ metrics [97]. For each replicate, each iBAQ value was normalized by the summed values of the 11 proteins. Error bars represent +/− SD. (B) and (C) Western blot validations in Benzonase treated and streptactin-purified and extracts. (D) HBc was immune-precipitated from nuclear extracts of dHepaRG-HBc (HBc), dHepaRG-ST-HBc (ST-HBc) and control dHepaRG (RG) cells induced with Tet for two days. Eluted protein were analyzed by western blot using anti HBc and anti-SRSF10 antibodies. The asterisk indicates the positions of IgG heavy chain.

### SRSF10 modulates HBV RNA levels

SRSF10, the most abundant cellular protein in our HBc nuclear interactome study, is a member of the SR protein family of splicing factors [30, 31]. Some of these proteins are uniquely nuclear-localized whereas others, such as SRSF10, can shuttle between the nucleus and the cytoplasm [32]. Analyses performed in HBV-infected primary human hepatocytes (PHH) indicated that, in this model, HBV infection did not alter the amount of this protein, which was mostly nuclear-localized (S2A-C Fig). The interaction between HBc and SRSF10 in HBV-infected PHH was further investigated using a proximity ligation assay (PLA) that permits detection of protein-protein interactions *in situ* (at distances <40nm) at endogenous protein levels. The presence of multiple signals in the nucleus of infected cells indicated that HBc and SRSF10 were in close proximity (S2D Fig). Similar results were obtained for RBMX, another important RBP found in ST-HBc complexes for which positive PLA signals were equally observed in the nucleus of HBV-infected PHH (S3 Fig). Altogether, these results strongly suggested that the interaction between HBc and these two RBPs was maintained in HBV-infected PHH without, however, inducing a major modification in their intracellular distribution.

We then investigated the effect of a SRSF10 knock-down (KD) on HBV infection. Optimization of the siRNA transfection protocol led to a significant level of protein KD in PHH without affecting NTCP levels, strongly suggesting that HBV internalization was not affected (Fig 4A-C). In PHH, KD of SRSF10 resulted in a significant increased accumulation of total HBV RNAs and pgRNA without affecting cccDNA level (Fig 4D). Similar results were observed in dHepaRG cells (S4A-D Fig). In sharp contrast, KD of RBMX resulted in opposite effects on HBV replication, with a decrease of all viral parameters, including cccDNA (Fig 4E-F, S4 Fig). These results indicated that RBPs found associated with HBc play distinct roles in the HBV life cycle. To determine whether SRSF10 KD had similar effect on an already established HBV infection, siRNA-mediated KD was also performed 7 days after the onset of infection, when replication has reached a plateau [33, 34]. In dHepaRG cells, a reproducible increase of HBV RNA could be observed following SRSF10 KD even if at a lower level as compared to cells in which the KD was performed before infection (S5 Fig). Altogether these results indicate that SRSF10 behaves as a restriction factor that modulates HBV RNA levels.

**Fig 4.**
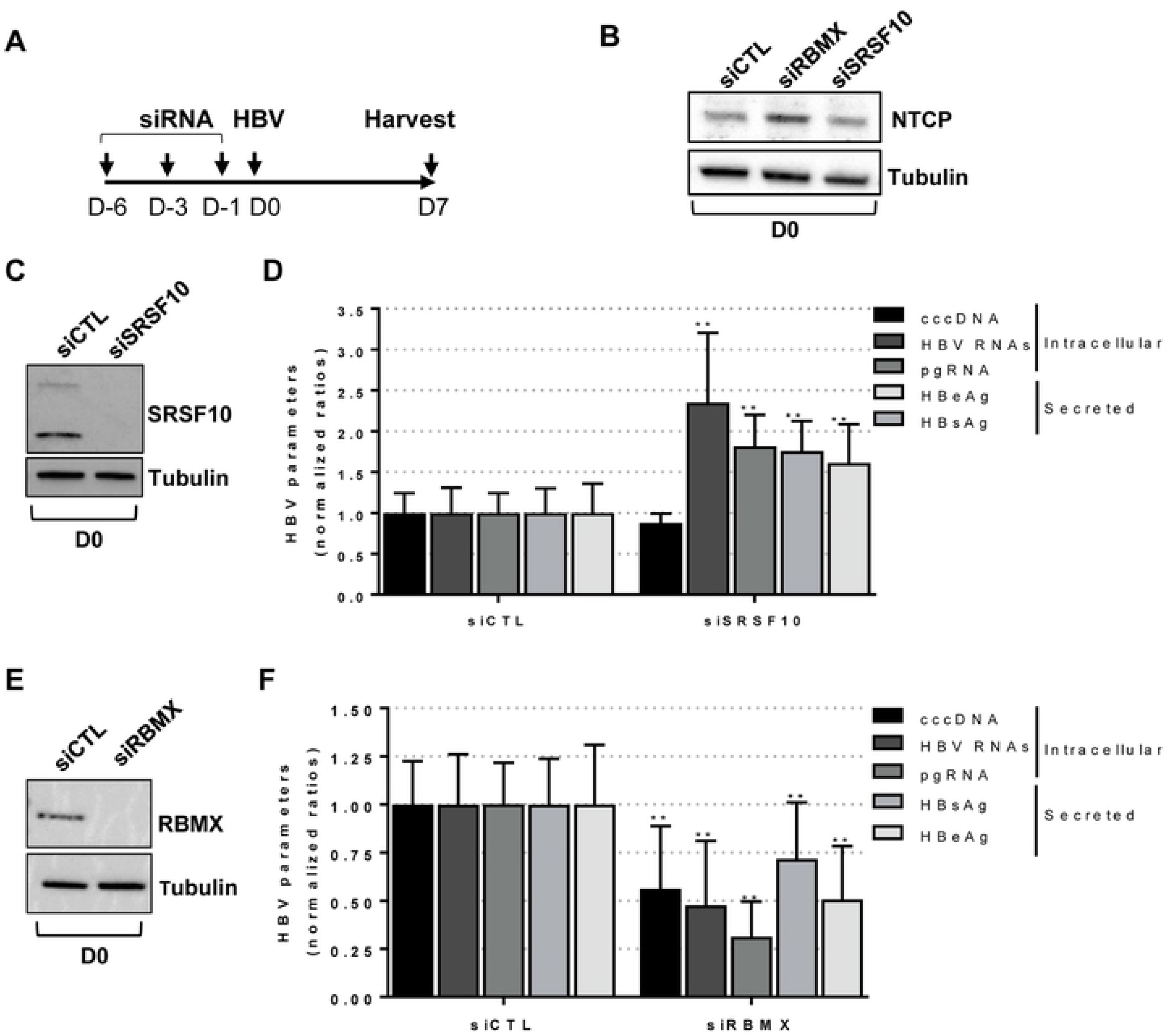
Effect of SRSF10 or RBMX KD on HBV replication in PHH. (A) Outline of the experimental protocol: cells were transfected with siRNA targeting SRSF10 or RBMX or control siRNA (siCTL) and then infected with HBV (MOI of 100 vge/cell). Cells and supernatants were harvested 7 days (D) post-infection (pi) and analyzed to measure intracellular and secreted HBV parameters. (B), (C) and (E) Western bot analysis of NTCP, SRSF10 and RBMX protein levels at D0 of the protocol, respectively. (D) and (F) HBV intracellular and extracellular parameters were measured at D7 pi. Results are expressed as the mean normalized ratio +/− SD, between siSRSF10 or siRBMX and siCTL transfected cells, of 3 independent experiments, each performed in triplicate, with PHH from different donors.

### A small molecule inhibitor of SRSF10 phosphorylation strongly impairs HBV replication and antigen secretion

SRSF10 activity was previously shown to be tightly controlled by phosphorylation, which regulates its interaction with other RBPs and splicing activities [35–38]. De-phosphorylation of SRSF10 occurs in response to heat shocks, DNA damage or during mitosis. More recently, compound 1C8 (Fig 5A), was shown to prevent SRSF10 phosphorylation, in particular at serine 133,in the absence of any other detectable effect on other SR proteins, and to inhibit HIV-1 replication with a combined effect on HIV-1 transcription and splice site selection likely producing an imbalance in viral protein required for replication [39, 40].

**Fig 5.**
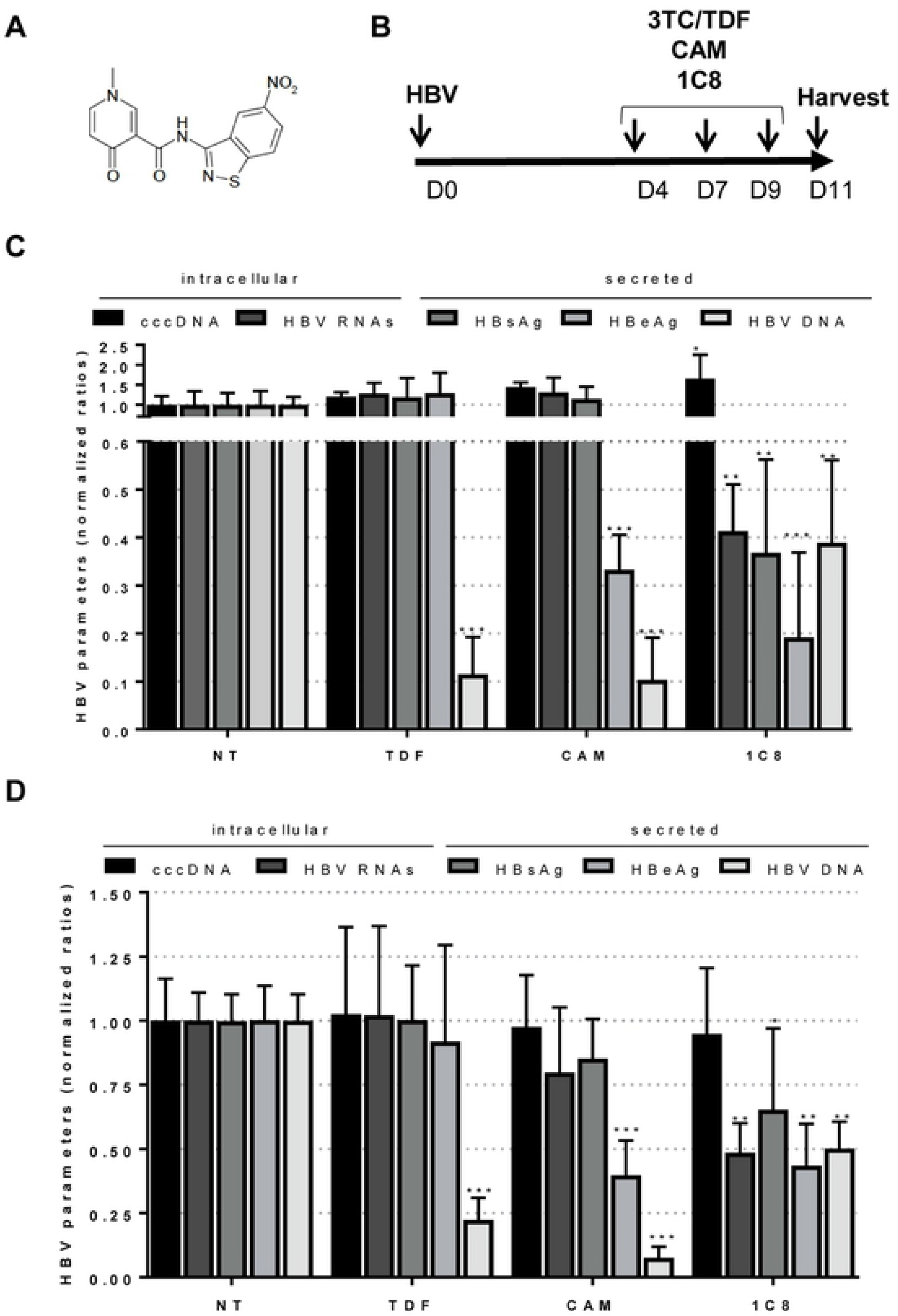
Effect of 1C8 on an established HBV infection. (A) Molecular structure of 1C8. (B) Outline of the experimental protocol: HBV-infected dHepaRG cells (**C**) or PHH (**D**) were treated three times with Tenofovir (TDF at 10μM), a Core allosteric modulator (CAM at 10μM) or 1C8 (10 μM) starting at D4pi. Intracellular and secreted HBV parameters were quantified 2 days after the last treatment. Results are expressed as the mean normalized ratio +/− SD between non-treated and treated cells of 3 independent experiments, each performed in triplicate.

To explore the effect of 1C8 on HBV replication we first assessed its effects when added on HBV-infected dHepaRG cells (Fig 5B). In this setting, treatment with 1C8 resulted in a strong decrease of viral RNAs and all downstream secreted parameters (Fig 5C). Interestingly, this phenotype was different from that observed with other antiviral compounds such as a NUC (Tenofovir) that uniquely inhibited HBV DNA synthesis, or a Core allosteric modulator (CAM) that additionally inhibited HBe secretion [41]. This effect, although weaker, was maintained in HBV-infected PHH (Fig 5D), a more relevant/physiologic model to assess the activity of compounds targeting host functions. Dose response analyses in dHepaRG indicated an effective concentration 50% (EC50) of approximately 10 and 5 μM for HBV RNAs/secreted DNA and HBsAg/HBeAg, respectively, in the absence of detectable cell cytotoxicity (S6 Fig). In subsequent analyses we sought to determine if 1C8 was equally active on other HBV genotypes than D that was used in all our previous experiments. In dHepaRG cells we found that 1C8 could inhibit the replication of HBV genotype C, with a significant decrease of viral RNAs and all secreted parameters suggesting that its antiviral effect is pan-genotypic (S7 Fig).

Altogether, these results indicate that 1C8 can inhibit HBV replication by reducing HBV RNA levels. The inhibitory effect of 1C8 on HBV RNAs, opposite to that observed following SRSF10 KD, suggests that the de-phosphorylated form of SRSF10, that is depleted following siRNA transfection and, in contrast, induced after 1C8 treatment, is responsible for the observed antiviral activities of this cellular RBP.

### 1C8 antiviral effect is partially dependent on SRSF10, promoting a reduction in HBV RNAs but not their splicing

To verify if the effect of 1C8 on HBV RNA accumulation was indeed related to SRSF10, experiments combining SRSF10 KD and 1C8 treatment were conducted (Fig 6A). Based on the model proposed, the inhibitory effect of 1C8 on viral RNA production, should be prevented by depleting SRSF10. As previously observed, each treatment alone, 1C8 or siSRSF10, resulted in opposite effects on HBV RNA levels. Remarkably, in cells receiving both treatments, depletion of SRSF10 could partially rescue the inhibitory effect of 1C8 to a level similar to that observed in control cells without, however, reaching that measured in siSRSF10-transfected cells (Fig 6B-C). This result indicates that the antiviral effect of 1C8 is dependent on SRSF10. The lack of a complete rescue, in cells treated with 1C8 and depleted of SRSF10, could be explained by the persistence of a low level of dephosphorylated SRSF10. Alternatively, it is also possible that 1C8 additionally targets other cellular and/viral factors that are involved in the anti-viral effect.

**Fig 6.**
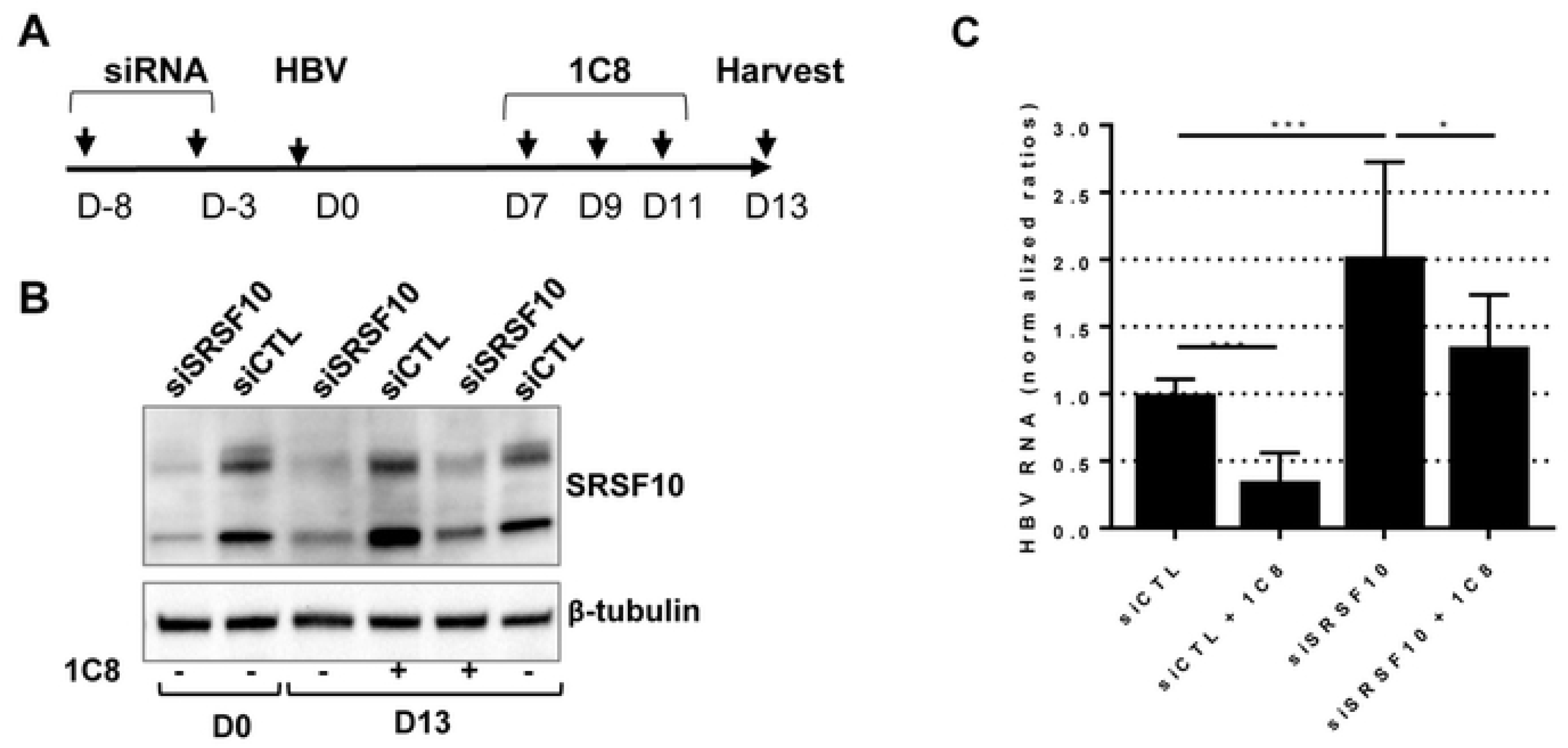
Combined effect of sRSF10 KD and 1C8 treatment on HBV-infected dHepaRG cells. (A) Outline of the experimental procedure: dHepaRG cells were transfected once or twice with siRNA targeting SRSF10, then infected with HBV (MOI of 250 vge/cell), and 7 days later treated three times with 1C8 (10μM). (B) Western blot validation showing SRSF10 depletion. (C) Quantification of intracellular HBV RNAs. Results are expressed as the mean normalized ratio +/− SD between treated and/or siSRSF10 transfected cells and siCTL-transfected cells of 3 independent experiments, each performed in triplicate.

All HBV RNAs required and sufficient for a productive replication (*i.e.* production of virion and viral proteins) are unspliced. Nonetheless, several spliced HBV mRNA have been documented in experimental models and more importantly in patient samples, indicating that, if an active mechanism of escape from splicing exists, it must be partial and/or ineffective at a certain stage during chronic infection [10]. Among the numerous HBV spliced RNAs, two major spliced forms result in the production of new viral proteins, some being potentially involved in viral pathogenesis, and particles containing shorter viral genomes [10, 42, 43]. In our previous assays, the primers used to quantify HBV RNAs (total and pgRNA) localized to an unspliced region of the HBV genome. Therefore it was possible that the variations in HBV RNA levels observed after SRSF10 KD or 1C8 treatment could be due to a specific modulation in some spliced variants or to a differential effect on spliced versus unspliced forms. To explore this possibility RNA extracted from siRNA transfected hepatocytes were analyzed by RT-qPCR using primers able to specifically detect each spliced and unspliced RNA (S2 Table). Unexpectedly, the relative quantification of each RNA variant in SRSF10-depleted versus control cells indicated that, both in dHepaRG and PHH, KD of SRSF10 resulted in a global increase of all HBV RNA variants, including all detected spliced forms without inducing a preferential modulation of a spliced versus unspliced variants (S9A-B Fig). Similarly, treatment of HBV-infected dHepaRG cells with 1C8 post-infection resulted in a strong reduction of all viral RNA whether spliced or unspliced (S9C Fig). These results indicated that the respective proviral or /antiviral effects of SRSF10 KD or 1C8 was not associated to a variation in the level of spliced versus unspliced HBV RNAs. They also suggested that both treatments acted on HBV RNAs synthesis and/or stability. To verify this point, total and nascent HBV RNAs were quantified following SRSF10 KD or 1C8 treatment. The quantification of nascent HBV RNAs was performed by labeling newly transcribed RNAs with ethynyl uridine (EU) for 2 hours before capture (Fig 7A and 7B). As expected, depletion of SRSF10 in dHepaRG cells prior to HBV infection increased total HBV RNAs. Actinomycin D (ActD), a global transcription inhibitor, strongly reduced the level of nascent RNA. In contrast, in cells transfected with siSRSF10, newly transcribed HBV RNAs were increased at a level similar to that observed for total RNAs (Fig 7C). The same analysis performed on 1C8-treated cells indicated that the compound equally reduced total and nascent HBV RNAs (Fig 7D). Altogether, these analyses indicate that SRSF10 and 1C8 did not modify the splicing level of HBV RNAs but, rather, that both treatments exert their effect by modifying the transcription and/or the stability of nascent viral RNAs.

**Fig 7.**
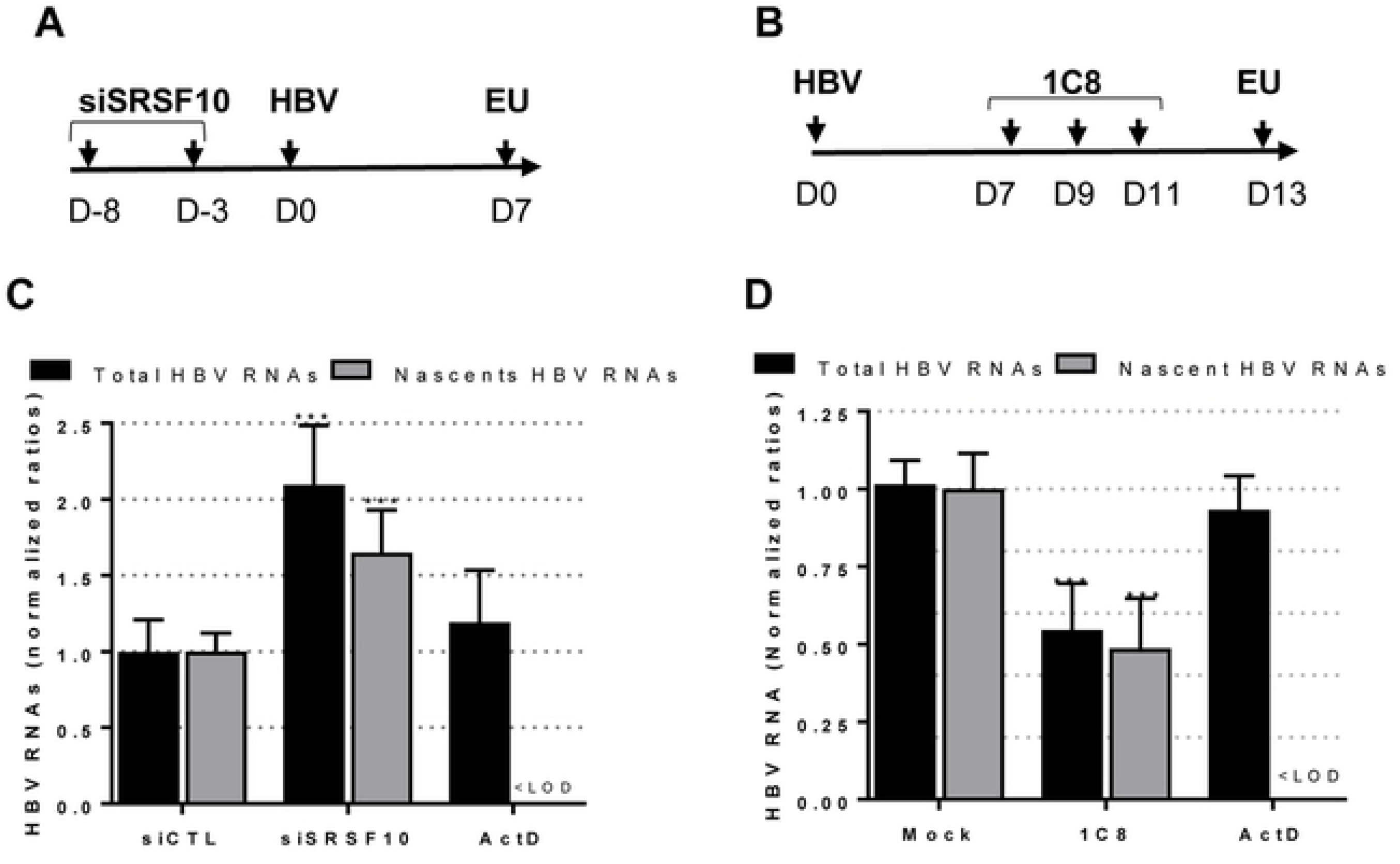
Analysis of nascent HBV RNAs following SRSF10 KD or 1C8 treatment. (A) dHepaRG cells were transfected with siRNA against SRSF10 and then infected with HBV (MOI of 250 vge/cell). Edu labelling was performed at D7pi for 2 hours. (B) dHepaRG cells were infected with HBV and then treated three times with 1C8 (40μM) at D7, D9 and D11pi. EU incorporation was performed at D13pi for 2 hours. (C) and (D) Run-on analyses. Intracellular RNA was extracted from transfected/treated cell cells and either directly quantified using HBV primers (Total HBV RNAs) or purified using the Click-iT Nascent RNA Capture kit to quantify newly synthetized RNAs (nascent HBV RNAs). Control was provided by treating cells with Actinomycin D (ActD at 10mg/ml) added to cells 20 min before labeling (see Methods). <LOD: under the limit of detection.

## DISCUSSION

Our strategy to expand the knowledge on HBc nuclear functions was based on the identification of its host protein partners in the nucleus of dHepaRG cells, when expressed alone in the absence of other viral constituents. HepaRG cells represented one of the best alternative models to freshly isolated human hepatocytes, and could be easily engineered as opposed to PHH [44]. This analysis indicated that in the nucleus HBc interacts with RBPs that are mainly involved in all steps of mRNA metabolism (Fig 2). Most of these RBPs localize in well-defined nuclear bodies, in particular speckles and paraspeckles [45, 46]. Recent studies have highlighted that many active genes are located in proximity to nuclear speckles and that this association results in an increase of nascent transcript levels [47, 48]. The association of HBc with cellular RBPs strongly suggests that it may intervene in the metabolism of viral RNAs by interacting with a highly interconnected network of proteins that can act at several transcriptional and post-transcriptional steps [49]. Accordingly, HBc has several features similar to those found in many cellular RBPs with, notably, a positively-charged CTD composed by a long stretch of arginines separated by seven serine residues resembling the arginine/serine-rich domain (RS domain) of several RBPs in particular of SR proteins [17, 50, 51]. When expressed in bacteria, HBc, thanks to its CTD, displays a strong RNA-binding activity [52]. Interestingly, several studies have shown that HBc also has a strong affinity for DNA, and can associate with cccDNA both *in vitro* and *in vivo* [24–27]. Although no evidence yet support the binding of HBc to viral and/or cellular RNA in the nucleus of HBV-infected hepatocytes, HBc, like some RBPs, may possess the dual ability to bind to DNA and RNA [53]. This activity might be tightly modulated by the phosphorylation level of its CTD, as previously demonstrated during pgRNA packaging [17]. Alternatively the association for HBc with cccDNA and/or RNA may be indirectly mediated *via* RNA molecules and/or its interaction with cellular RBPs. Interestingly, some RBPs found associated to HBc, such as RBMX (Fig 1) were also reported to bind to DNA, in particular during DNA repair [54, 55].

Among the RBPs interacting with HBc, we focused our studies on SRSF10, a member of the SR proteins family that was the most highly abundant in HBc-containing complexes. As all the other members of the SR family, SRSF10 is composed of a N-terminal RNA recognition motif (RRM) and a C-terminal RS domain that is responsible for binding to other RBPs [32]. Two isoforms of SRSF10 of 37 and 20 KDa have been described, the smaller presenting a deletion of the C-terminal domain, but only the full-length has been extensively studied. Initial studies identified SRSF10 as a splicing repressor when de-phosphorylated in response to heat shock [38, 56]. Further analyses showed that SRSF10 is a regulator of AS, depending on its phosphorylation level that also determines its interaction with diverse RBPs, in particular TRA2A, TRA2B, hnRNPK, F, and H [35, 38, 57]. Importantly, SRSF10 influences the AS of several cellular transcripts involved in pathways of stress response, DNA damage response, apoptosis, and carcinogenesis [57–59]. In accordance to its role in the stress response, SRSF10 was described as a component of paraspeckles, nuclear stress bodies and cytoplasmic stress granules [60–62]. SRSF10 also plays a role in the control of viral RNA, in particular of HIV-1 [40, 63].

To investigate the role of SRSF10 during HBV replication we analyzed the consequences of its KD in differentiated hepatocytes. Depletion of SRSF10 induced a reproducible increase in viral RNAs, proteins and secreted DNA that was observed in both dHepaRG and PHH (Fig 4 and S4 Fig). These results suggest that SRSF10 may normally repress the production of viral RNAs. The importance of SRSF10 in the HBV life cycle was further suggested by the finding that compound 1C8, previously characterized as an inhibitor of SRSF10 phosphorylation [40], strongly inhibited HBV replication by inducing a marked reduction of viral RNA levels in both HBV-infected dHepaRG cells and PHH (Fig 5). The opposite effects observed upon SRSF10 depletion and 1C8 treatment strongly suggest that the de-phosphorylated form of SRSF10 is responsible for the restriction effect. According to this hypothesis, while treatment with 1C8 results in the accumulation of de-phosphorylated SRSF10, leading to a strong inhibitory phenotype, depletion of SRSF10 using siRNA affects all SRSF10 forms whether phosphorylated or not, thereby attenuating its restrictive activity. The involvement of SRSF10 in 1C8 antiviral effect is further suggested by the finding that it could be partially reverted upon SRSF10 depletion. As previously shown during AS, de-phosphorylation of SRSF10 may change its interaction with other cellular proteins important for HBV life cycle as well its capacity to bind viral RNA [38, 40]. It is also possible that 1C8 prevents the phosphorylation of other targets in addition to SRSF10. In particular, 1C8 may interfere with the phosphorylation of HBc in the nucleus, with consequences on its capacity to bind not only to other factors, in particular RBPs, but also DNA and/or RNA. Phosphorylation of HBc at serine residues within its CTD has been described as an important event to regulate pgRNA packaging and reverse transcription during nucleocapsid formation in the cytoplasm [17, 64–68]. An attractive hypothesis is that phosphorylation of the nuclear pool of HBc may similarly control its capacity to bind to DNA and/or RNA by modulating its positive charges, as previously demonstrated by studies performed in bacteria [52]. The identification of kinase(s) inhibited by 1C8 will be critical to further decipher its mode of action and identify other potential target(s).

Finally, one surprising finding of our study is that neither SRSF10 nor 1C8 altered the splicing of HBV RNAs, as compared to what observed for other viruses (S9 Fig) [40, 69]. While this result does not exclude that part of their effects on HBV may be indirectly mediated by an alteration in splicing of other cellular RNAs, our data rather suggest that both SRSF10 and 1C8 may act by controlling the level of nascent HBV RNAs (Fig 7). Beside their canonical role during splicing, several SR proteins were found to act at several other steps of cellular and viral RNA metabolism including transcription, stability and nuclear egress [31, 70, 71]. Notably, besides pre-mRNA splicing, some SR proteins have been reported to directly or indirectly associate to the phosphorylated CTD of RNA polymerase II and to stimulate transcriptional elongation [72, 73]. Interestingly, several RBPs previously identified as important for HBV replication were shown to participate in the control of HBV transcription and/or in RNA stability. This is the case in particular for the splicing factors HNRNPC, HNRNPK, PUF60, RBM24 and TARBP [74–78]. It is therefore conceivable that SRSF10 may differentially associate with HBV nascent RNA in the nucleus to control its synthesis/elongation and/or its stability. Recent studies have indicated that HBV RNA stability is controlled by m-6A methylation [79, 80]. SRSF10 may intervene in HBV RNA stability by interacting with m-6A readers or interfering with their RNA binding activity [81]. Future studies analyzing epigenetic changes and modifications in HBc/SRSF10 cccDNA/RNA binding activities in the presence of 1C8 should help uncover the underlying mechanisms.

Altogether, our study revealed that nuclear HBc can connect to an array of cellular RBPs. It also identified SRSF10 as a restriction factor in HBV viral RNA production, therefore providing a basis for the evaluation of a new class of host-targeted antiviral compounds that could improve the current anti-HBV arsenal and encourage combinational therapies.

## Methods

### Cell culture and HBV infection

HepaRG cells were cultured, differentiated, and infected by HBV as previously described [33]. HepaRG-TR-HBc and HepaRG-TR-ST-HBc were obtained by transducing HepaRG-TR cells [82] with Lenti4/TO lentiviral vectors expressing either HBc or ST-HBc under the control of the minimal CMV/TetOn promoter. Details on sequences are available upon request. Transduced cells were selected using blasticin (10 μg/mL) and zeocyn (100 μg/mL), then amplified and frozen as polyclonal lines. Primary human hepatocytes (PHH) were freshly prepared from human liver resection obtained from the Centre Léon Bérard (Lyon) with French ministerial authorizations (AC 2013-1871, DC 2013 – 1870, AFNOR NF 96 900 sept 2011) as previously described [83]. HBV genotype D inoculum (subtype ayw) was prepared from HepAD38 [84] cell supernatant by polyethylene-glycol-MW-8000 (PEG8000, SIGMA) precipitation (8% final) as previously described [85]. HBV genotype C viral inoculum was similarly prepared from the supernatant of a newly developed stably-transformed HepG2 cell line. Briefly, the cell line was obtained by transfection of a linearized pcDNA3Neo-HBV plasmid containing 1.35 genome unit of a consensus sequence of HBV genotype C (obtained from HBV database: https://hbvdb.lyon.inserm.fr/; ref [86]) and a double-round selection under G418 (500 ng/mL) by colony cell cloning (very low density seeding in large flasks). The titer of endotoxin free viral stocks was determined by qPCR. Cells were infected overnight in a media supplemented with 4% final of PEG, as previously described [33]. Infection was verified by measuring secreted HBsAg and HBeAg 7 to 10 days later by CLIA (Chemo-Luminescent Immune Assay) following manufacturer’s instructions (AutoBio, China).

### Chemical reagents

Unless otherwise specified, chemical reagents, drugs, antibiotics were purchased from Sigma Aldrich. Tenofovir (TDF) was a kind gift of Gilead Sciences (Foster city, USA). The core assembly modulator (CAM) used in the experiments was previously described [41], and resynthesized by AI-Biopharma (Montpellier, France). 1C8, *i.e.* 1C8, *i.e.* 1-Methyl-N-(5-nitrobenzo[d]isothiazol-3-yl)-4-oxo-1,4-dihydropyridine-3-carboxamide, was synthesized in Dr Gierson laboratory, but also re-synthesized and purified at 99% by AGV (Montpellier, France).

### Purification of ST-HBc complexes and sample preparation for mass spectrometry

HepaRG-TR-HBc and HepaRG-TR-ST-HBc cells (1,25×10^8^ cells) were first differentiated, then transgene expression induced for 72 hours by adding Tet (5 μg/ml) in the culture media. Nuclei were purified from cells using the NE-PERT^M^ Nuclear and Cytoplasmic Extraction kit (Thermo Scientific), and nuclear extracts prepared by suspending nuclei in NER solution in the presence of protease inhibitors (Complete EDTA-free Protease Inhibitor Cocktail, Roche) and digestion or not with Benzonase (Sigma-Aldrich), during 40 min on ice. Nuclear extract recovered after centrifugation were added on Strep-Tactin^R^ gravity columns (IBA) and further purified following manufacturer’s instructions.

### Capsid migration assay

The intracellular formation of HBV nucleocapsids was assessed by native agarose gel electrophoresis of cell lysates, followed by transfer onto the enhanced chemiluminescence membrane and western blot analysis, as described previously [87].

### Mass spectrometry-based quantitative proteomic analyses

Eluted proteins were stacked in a single band in the top of a SDS-PAGE gel (4-12% NuPAGE, Life Technologies) and stained with Coomassie blue R-250 before in-gel digestion using modified trypsin (Promega, sequencing grade) as previously described [88]. Resulting peptides were analyzed by online nanoliquid chromatography coupled to tandem MS (UltiMate 3000 and LTQ-Orbitrap Velos Pro, Thermo Scientific). Peptides were sampled on a 300 μm x 5 mm PepMap C18 precolumn and separated on a 75 μm x 250 mm C18 column (PepMap, Thermo Scientific) using a 120-min gradient. MS and MS/MS data were acquired using Xcalibur (Thermo Scientific).

Peptides and proteins were identified and quantified using MaxQuant (version 1.5.3.30) [89] using the Uniprot database (*Homo sapiens* taxonomy proteome, October 2016 version), the sequence of HBc, and the frequently observed contaminant database embedded in MaxQuant. Trypsin was chosen as the enzyme and 2 missed cleavages were allowed. Peptide modifications allowed during the search were: carbamidomethylation (C, fixed), acetyl (Protein N-ter, variable) and oxidation (M, variable). Minimum peptide length was set to 7 amino acids. Minimum number of peptides, razor + unique peptides and unique peptides were set to 1. Maximum false discovery rates − calculated by employing a reverse database strategy − were set to 0.01 at PSM and protein levels. The “match between runs” option was activated. The iBAQ value [90] calculated by MaxQuant using razor + unique peptides was used to quantify proteins.

Statistical analysis were performed using ProStaR [91]. Proteins identified in the reverse and contaminant databases and proteins exhibiting less than 3 quantification values in one condition were discarded from the list. After log2 transformation, iBAQ values were normalized by median centering before missing value imputation (2.5-percentile value of each sample). Statistical testing was conducted using *limma* test. Differentially-expressed proteins were sorted out using a log_2_(fold change) cut-off of 2 and a p-value cut-off allowing to reach a FDR inferior to 1% according to the Benjamini-Hochberg procedure (p<0.0079 and p<0.0063 for dataset with and without Benzonase, respectively).

### Gene Ontology analyses and construction of the HBc-interactome

Proteomics data from the two conditions (without (benz-) or with (benz+) benzonase) were filtered out according to p-value (<0.005) and fold change (>4). Significant proteins common to the two conditions were extracted with a Venn diagram using their UniProtKB accession numbers. Statistical overrepresentation tests of these proteins were computed with PantherDB 11.1 and GO complete annotation sets. Overrepresented protein accession numbers were selected to further build and analyze their interacting network by means of Cytoscape software 3.5.1, querying IntAct molecular interaction database (May 27, 2017) with PSICQUIC service application 3.3.1, and Network Analyzer application 3.3.2.

### siRNA transfection

dHepaRG or PHH cells seeded into a 24-well plate were transfected with 25 nM or 10 nM of siRNAusing Dharmafect#1 (GE HealthCare) or Lipofectamine RNAiMax (Life Technologies), respectively, following manufacturer’s instructions. SiRNA used were the following: siSRSF10 (Dharmacon SmartPool L-190401), siRBMX (Dharmacon SmartPool L-011691), siControl (Dharmacon D-001810).

### Co-immunoprecipitation and western blot analysis

For Co-IP analyses, 300-500 μg of nuclear extracts, prepared as indicated above, were precleared with Protein A/G magnetic beads (Pierce™) for 2hrs at 40°C and then incubated over-night at 4°C on a rotating wheel with 2 μg of anti HBc antibody (Dako B0586). Immune-complexes were captured with protein A/G magnetic beads, washed four times in IP buffer and then eluted by boiling for 5 min in 2X loading buffer (Laemmli). For western blot, proteins were resolved by SDS-PAGE and then transferred onto a nitrocellulose membrane. Membranes were incubated with the primary antibodies corresponding to the indicated proteins. Proteins were revealed by chemi-luminescence (Super Signal West Dura Substrate, Pierce) using a secondary peroxydase-conjugated antibody (Dako) at a dilution of 1:10000. Primary antibodies used were: anti–HBc (Ab140243, 1/1000) or a home generated anti-HBc (1/40000; generous gift from Dr Adam Zlotnick, Bloomington, USA), anti-SRSF10 (Ab77209, 1/2000), anti-RBMX (Ab190352, 1/2000), anti-TRA2B (Ab171082, 1/2000), anti-SRSF1 (Ab38017, 1/1000), anti-SRSF2 (Ab204916, 1/1000), anti –DDX17 (Proteintech 19910-1-AP, 1/1000), anti-PARP1 (Ab6079, 1/1000), anti-DNAJB6 (Ab198995, 1/1000), anti-β-Tubulin (Ab6044, 1/10000), anti-Lamin B1 (Ab16048, 1/10000).

### Immunofluorescence analyses

Analyses were performed as described previously using Alexa Fluor 488 or 555 secondary antibodies (Molecular Probes) [88]. Primary antibodies used were: anti-HBc (Thermo MA1-7607, 1/500), anti-SRSF10 (SIGMA HPA053831, 1/50), anti-RBMX (Ab19035, 1/500). Nuclei were stained with Hoescht 33258. To get rid of the auto fluorescence of PHH, the images were collected on a confocal NLO-LSM 880 microscope (Zeiss) by spectral imaging and linear unmixing with the online finger-printing acquisition mode of ZEN Black software. To achieve the linear unmixing, the spectrum of each anti-body coupled fluorochromes and Hoescht was recorded individually on non-auto fluorescent dHepaRG cells. The auto fluorescence spectrum was acquired for each PHH batch using unlabeled cells. Further image processing was performed using ICY [92]. Each experiment was reproduced several times, and the images are representative of the overall effects observed under each condition.

### Nucleic acid extractions, reverse transcription and qPCR analyses

Total RNA and DNA were extracted from cells with the NucleoSpin RNA II and Nucleospin® 96 tissue kit, respectively, according to the manufacturer’s instructions (Macherey-Nagel). RNA reverse transcription was performed using SuperScript III (Invitrogen). Quantitative PCR for HBV were performed using HBV specific primers and normalized to PRNP housekeeping gene as previously described [93]. Pre-genomic RNA was quantified using the TaqMan Fast Advanced Master Mix (Life Technologies) and normalized to GusB cDNA levels. HBV cccDNA was quantified from total DNA following digestion for 45 min at 37°C with T5 exonuclease (Epicentre) to remove rcDNA followed by 30 min heat inactivation. cccDNA amount was quantified by TaqMan qPCR analyses and normalized to β-globin cDNA level, as previously described [94].

### Analysis of spliced HBV RNA

The analysis of HBV spliced RNA was performed the RNomics platform of the University of Sherbrooke (Canada) as previously described [95, 96]. After reverse-transcription, quantitative qPCR was performed using primers designed to detect each spliced and unspliced RNA and normalized to the MLRP19, PUM1 et YWHAZ genes (S8 Table). Primers were designed to detect 15 spliced RNA (sv1 to sv15) and 3 intronic regions (intron 1, 2 and 2b), as described in ref. [10].

### Quantification of nascent HBV RNA

HBV nascent RNA were quantified using the Click-iT^®^ Nascent RNA Capture Kit (Life Technologies) following the manufacturer’s instructions. Briefly HBV-infected dHepaRG were incubated for 2 hours with 5-ethynyl Uridine (EU) before RNA extraction and biotinylation. Control was provided by cells treated with Actinomycin D (1 μM) 20 min before labeling. Biotinylated RNA was purified on streptavidin magnetic beads. Total and EU-labeled RNA was reverse-transcribed and quantified as indicated above.

### Viability/cytotoxicity assays

Viability/cytotoxicity was assessed using the CellTiter-Glo^R^ Luminiscent assay (Promega) following the manufacturer’s instructions.

### Statistical analysis

Statistical analyses were performed using the XLStat software and Kruskal-Wallis tests with multiple comparison respect to non-treated cells (Dunn’s post-test). For all tests, a p value ≤ 0,05 was considered as significant. * correspond to p value ≤ 0.05; ** correspond to p value ≤ 0.01; *** correspond to p value ≤ 0.001.

## Acknowledgments

We would like to thank Adam Zlotnick for providing the anti-HBc antibody, Christophe Vanbelle (Imaging platform of CRCL) for his help on confocal microscope analyses, and Brieux Chardès and Claire Bugnot for technical assistance. We also would like to thank Laura Dimier, Jennifer Molle, Océane Floriot and Anaëlle Dubois for their help with the isolation of primary human hepatocytes, as well as the staff from Pr. Michel Rivoire’s surgery room for providing us with liver resection. We also thank Philippe Thibault, Elvy Lapointe and Mathieu Durand at the RNA platform of Université de Sherbrooke.

## Supporting information

**S1 Fig. Functional analysis of dHepaRG-TR-ST-HBc cells**. (A) Immunofluorescence (IF) of analysis of HBc localization in dHepaRG-TR-ST-HBc versus HBV-infected dHepaRG and PHH. (B) Intracellular HBV capsids, produced by the indicated cell lines, were analyzed by native gel electrophoresis followed by western blot with anti-HBc antibody. Lanes: 1.HepG2.2.15 2.dHepaRG-TR-HBe; 3.dHepaRG-TR-HBc; 4.dHepaRG-TR-ST-HBc.

**S2 Fig. Intra-cellular distribution of SRSF10 in HBV-infected hepatocytes and co-localization with HBc**. (A) PHH were infected with HBV (MOI of 500 vge/cell) or mock-infected (NI) and harvested at different days pi. Productive infection was verified by quantification of HBsAg and HBeAg levels in the supernatant of infected cells at D7 and D9pi. (B) Nuclear (N) and cytoplasmic (C) extracts were prepared and analyzed by western blot using anti-SRSF10 and anti-HBc antibodies. Cell fractionation was analyzed using anti-Lamin B1 and anti-β-Tubulin antibodies. (C) and (D). HBV-infected PHH were analyzed at D9pi by either immunofluorescence (C) or PLA (D) using anti SRSF10 and anti-HBc antibodies. Images were taken using a Zeiss 880 confocal microscope. Scale bar: 10μm.

**S3 Fig. Intra-cellular distribution of RBMX in HBV-infected hepatocytes and co-localization with HBc**. PHH were infected with HBV (MOI of 500 vge/cell) or mock-infected (NI) and analyzed at D9pi by either immunofluorescence (A) or PLA (B) using anti-RBMX and anti-HBc antibodies. Images were taken using a Zeiss 880 confocal microscope. Scale bar: 10μm.

**S4 Fig. Effect of SRSF10 or RBMX KD on HBV replication in dHepaRG cells**. (A) Outline of the experimental protocol in dHepaRG cells: cells were transfected with siRNA targeting SRSF10 or RBMX or control siRNA (siCTL) and then infected with HBV (MOI of 250 vge/cell). (B) NTCP levels in siRNA transfected dHepaRG cells before HBV infection (D0). **C.** Western blot validations in cells secreted parameters measured at D7pi. Results are expressed as the mean normalized ratio +/− SD, between siSRSF10 or siRBMX and siCTL transfected cells, of 3 independent experiments, each performed in triplicate.

**S5 Fig. Effect of SRSF10 on established HBV replication**. (A) Outline of the experimental protocol: dHepaRG cells were infected with HBV (MOI of 250 vge/cell) and then transfected twice with siRNA targeting SRSF10 or control siRNA (siCTL). Cells and supernatants were harvested at D15pi and analyzed to measure extracellular and intracellular HBV parameters. (B) Western blot validation of SRSF10 KD. (C) Effect of SRSF10 KD on intracellular and secreted HBV parameters. Results are expressed as the mean normalized ratio +/− SD, between siSRSF10 or siRBMX and siCTL transfected cells, of 3 independent experiments, each performed in triplicate.

**S6 Fig**. (A) to (D) **Measure of 1C8 EC50 on HBV-infected dHepaRG.** dHepaRG cells were infected with HBV (MOI of 250 vge/cell) for 7 days followed by three treatments with increasing concentration of 1C8. Total HBV RNAs (A), secreted HBV DNA (B), HBsAg (C) and HBeAg (D) were measured two days after the last treatment. Results are presented as the mean change in expression or secretion +/− SD of three independent experiments, each performed in triplicate. (E) **Toxicity assay.** Cell viability of dHepaRG cells treated with increasing concentrations of 1C8, was measured using the CellTiter-Glo® Luminiscent Cell Viability Assay (Promega). Non-infected (NI) and HBV-infected dHepaRG cells treated with DMSO and puromycin were used as negative and positive controls, respectively.

**S7 Fig. Effect of 1C8 on HBV (genotype C) replication in dHepaRG cells**. Cells were infected (MOI of 100 vge/cell) and treated as indicated in figure 5A. Treatments included, Tenofovir (TDF at 10 μM), a Core allosteric modulator (CAM at 10 μM) or 1C8 (10 μM). Intracellular and secreted HBV parameters were quantified 2 days after the last treatment. Results are expressed as the mean normalized ratio +/− SD between non-treated and treated cells of 2 independent experiments, each performed in triplicate.

**S8 Fig. Analysis of spliced and unspliced HBV RNA following SRSF10 KD or 1C8 treatment**. (A) and (B) Total RNA were extracted from dHepaRG (A) and PHH (B) transfected with siRNA following the previously described protocol (Fig 4A). (C) HBV-infected dHepaRG were treated with 1C8 as previously described (Fig 5B). HBV RNAs were analyzed by end-point RT-qPCR using sets of primers able to discriminate each spliced and unspliced form (see Materials and Methods section). Results are expressed as the mean ratio +/− SD between siSRSF10 and siCTL transfected cells of 3(A and C) or 2 (B) independent experiments.

**S1 Table. HBc interactome analysis using MS-based quantitative proteomics**. ST-HBc-associated proteins were identified using MS-based proteomics. For this, cell lysate expressing ST-HBc or untagged HBc were treated or not with Benzonase before purification by affinity using Strep-Tactin. Eluted proteins were digested with trypsin and the resulting peptides submitted to MS-based proteomic analysis. The proteins were then identified and quantified in each sample before statistical analysis allowing to sort out proteins enriched with ST-HBc compared to untagged HBc used as negative control.

**S2 Table. List of primers used for the quantification of spliced and unspliced HBV RNAs**.

